# Phylogenetic diversity of flowering plants declines across the growing season in Rocky Mountain wildflower communities

**DOI:** 10.1101/2023.11.06.565878

**Authors:** Leah N Veldhuisen, Brian J. Enquist, Katrina M Dlugosch

**Affiliations:** University of Arizona

**Keywords:** co-flowering, network analysis, Rocky Mountain Biological Laboratory, phylogenetic diversity, elevational gradient

## Abstract

Phylogenetic diversity is an important axis of biodiversity associated with many ecosystem functions, but its variation over time in temperate communities has rarely been explored. Plants’ flowering phenology is key for reproduction: it determines synchrony among plants, mutualists, antagonists, and favorable environmental conditions. Individual species’ flowering phenologies combine to determine which species’ flowers co-occur in time and space to form co-flowering communities. We recorded plants’ flowering phenology across elevations in the Colorado Rocky Mountains and predicted: 1) flowering species will form subsets of co-flowering species, 2) groups of co-flowering species will be more distantly related than the broader community, 3) marginal early and late season conditions will constrain flowering to species groups that are less phylogenetically diverse and 4) the entire community will have higher phylogenetic diversity at higher elevations. We identified co-flowering groups using a network analysis and quantified their phylogenetic diversity. Co-flowering communities had higher phylogenetic diversity early in the season and at higher elevations, but became less diverse by all metrics at the end of the season across elevations. Our results indicate that phylogenetic diversity varies consistently across the season and that conditions in the late season and at low elevations might constrain diversity.

## Introduction

How biological diversity is distributed through space and time has been a central question in ecology and evolution for centuries (Linnaeus 1799; Darwin 1859; MacArthur 1969; Hillebrand 2004; Schluter and Pennell 2017). Most recently, the distribution of evolutionary (phylogenetic) diversity across contemporary communities has received a great deal of attention (Webb et al. 2002; Mayfield and Levine 2010; Molina-Venegas et al. 2021; Tietje et al. 2023).

Higher phylogenetic diversity within a community indicates greater evolutionary divergence of community members, and it is thought that phylogenetic diversity may capture ecologically relevant variation among species that is missed when quantifying diversity by species richness or variation in one or a few traits (Cadotte et al. 2009; Owen et al. 2019; Huang et al. 2020; E-Vojtkó et al. 2023). Phylogenetic diversity is often observed to correlate poorly with variation in individual traits (CaraDonna and Inouye 2015; Gerhold et al. 2015; Maitner et al. 2022; E-Vojtkó et al. 2023), yet it does appear to be associated with community-level functions (Mazzochini et al. 2019; Molina-Venegas et al. 2021). For example, phylogenetic diversity correlates more positively with biomass than do other diversity metrics (Cadotte et al. 2008), and is associated with greater ecosystem stability and productivity (Cadotte et al. 2012; Cadotte 2013; Davies et al. 2016; Mazzochini et al. 2019) and increased resistance to species invasion (Galland et al. 2019). Extinction risk has also been observed to be phylogenetically clustered, indicating phylogenetic variation in the ecological vulnerability of species (Willis et al. 2008; Eiserhardt et al. 2015; González-Orozco et al. 2016). Here, we investigate patterns of phylogenetic diversity in plants that flower together in the same plant communities, and how their diversity changes across the growing season.

The timing of flowering (flowering phenology) is a key component of plant reproduction, as it determines the synchrony (or lack thereof) between plants and their reproductive mutualists and antagonists (Elzinga et al. 2007; Forrest and Miller-Rushing 2010; Munguía-Rosas et al. 2011; Freimuth et al. 2022), and with the environmental conditions favorable for reproduction (Jentsch et al. 2009). At the community level, individual species’ flowering phenologies combine to determine which species’ flowers co-occur in time and space (co-flower), shaping the floral resources available to pollinators (Blaauw and Isaacs 2014), and affecting whether plant species might compete with or facilitate one another to attract pollinators (Levin and Anderson 1970; Brown and Kodric-Brown 1979; Moeller 2004; Moeller and Geber 2005*a*; Ghazoul 2006; Kharouba et al. 2018). Importantly, nearly 90% of flowering plants rely on animal pollinators for reproduction (Ollerton et al. 2011), but pollinator populations are declining worldwide (Biesmeijer et al. 2006; Koh et al. 2016), disrupting plant-pollinator networks and driving declines in plant populations (Burkle et al. 2013; Potts et al. 2016). Plants are also changing their phenology in response to climate change and biodiversity loss (Menzel et al. 2006; Cleland et al. 2007; CaraDonna et al. 2014; Wolf et al. 2017), and such shifts in phenology are expected to alter interspecific interactions (Kharouba et al. 2018; Pareja-Bonilla et al. 2023; Austin et al. 2024). An understanding of how the phylogenetic composition of co-flowering plant communities changes across the growing season could provide a foundation for increasing our understanding of the relationship between species composition and functioning of these communities, and the potential consequences of environmental change and biodiversity loss for flowering.

Of particular importance is when in the season phylogenetic diversity peaks and when it is lowest, given the positive associations between phylogenetic diversity and ecosystem productivity and stability (Cadotte et al. 2012; Cadotte 2013; Davies et al. 2016; Mazzochini et al. 2019), and the potential sensitivity of low diversity communities to phylogenetically clustered declines or extinctions (Willis et al. 2008; Eiserhardt et al. 2015; González-Orozco et al. 2016). A central prediction of community phylogenetics is that ‘environmental filtering’ that favors only the species adapted to a particular set of conditions will result in a lower diversity of species establishing in a community, due to phylogenetic conservation of multi-trait strategies for the necessary environmental adaptations (Rejmánek 1996; Webb 2000; Webb et al. 2002). Evidence for this pattern has been found more often at broad spatial or environmental scales and environmental extremes, though not universally (Ma et al. 2016; Park et al. 2020; Maitner et al. 2022). For flowering communities, this would predict that groups of species co-flowering at one time of the season would be less diverse (more closely related) than the community as a whole, particularly at environmental extremes that might occur early and late in the season at the margins of suitable flowering conditions.

In contrast, closely-related species are predicted to compete more strongly at the local scale, leading to communities of more distantly related species (higher phylogenetic diversity), now known as the ‘competition-relatedness hypothesis’ (Darwin 1859; Ricciardi and Mottiar 2006). This hypothesis would predict that groups of species co-flowering and interacting would be more diverse than expected by chance than the community as a whole. Considering both hypotheses of environmental filtering and competition-relatedness, we might expect local-scale co-flowering communities to be overall more diverse than expected, but to become more closely-related and less diverse at the environmental extremes of the season, due to increased environmental filtering. Nevertheless, studies of community phylogenetic assembly have found mixed support for environmental filtering and competition-relatedness hypotheses (Daehler 2001; Procheş et al. 2007; Thuiller et al. 2010; Jones et al. 2013; Ma et al. 2016; Cadotte et al. 2018; Park et al. 2020). A persistent challenge has been addressing how the niche and fitness differences of potentially competing species relate to phylogeny, given that phylogeny is often not aligned with specific traits (Mayfield and Levine 2010; Maitner et al. 2022). Indeed, for flowering communities, floral traits are known to be highly variable phylogenetically (Vamosi and Vamosi 2010; CaraDonna and Inouye 2015; O’Meara et al. 2016), and so any phylogenetically structured environmental filtering or competitive interactions may be shaped by other phylogenetically conserved non-floral traits of species (but see (Martín-Hernanz et al. 2023; Xiang et al. 2023).

Despite the importance of understanding the diversity of co-flowering communities and testing predictions for their structure, little work has quantified the phylogenetic relationships of co-flowering species. Studies on floral networks by Albor et al. (2020, 2022) found low phylogenetic signal in the overlap of co-flowering species pairs but did not examine variation in phylogenetic diversity of co-flowering communities. Studies by Wolowski et al. (2017) of hummingbird-pollinated plants in the Brazilian Atlantic rainforest, and by Shrestha et al. (2019) of floral color communities in Australian herbs, each found some evidence for non-random phylogenetic structure (either less or more diverse than expected by chance), depending on the environment and subset of the community considered. Notably, Wolowski and colleagues found lower diversity and potential evidence of environmental filtering in lowland relative to montane sites, but they did not quantify phylogenetic diversity over time during the season (Wolowski et al. 2017). Shrestha and colleagues found variable temporal patterns in their data, depending on the pollinator type of the floral community analyzed (Shrestha et al. 2019). To our knowledge, no other studies have examined phylogenetic composition of flowering communities across a season.

Within a flowering community at a particular site, groups of species that flower at the same time within a season can be identified using the framework of a network analysis (Proulx et al. 2005; Arceo-Gómez et al. 2018). A network analysis quantifies the connections (‘edges’) between entities (‘nodes’) and identifies whether certain groups of nodes form clusters of strong connections to one another (a ‘module’), with weaker connections outside of the group. As proposed by Arceo-Gómez and colleagues (2018), the duration and intensity of flowering overlap can be used to define the strength of connections (edges) between pairs of flowering species (nodes), and these pairwise connections can be used to form a network of flowering interactions across the entire community. The extent to which the network has groups (modules) of species with more interactions with one another than expected by chance can be quantified as the network’s modularity. Modules then represent subsets of species within communities that co-flower more strongly with each other than with other species (Arceo-Gómez et al. 2018). Such network analyses have been used to investigate trait similarity within co-flowering modules and how plant-pollinator interactions may impact community assembly processes (Albor et al. 2020, 2022; Wei et al. 2021; Suárez-Mariño et al. 2022).

Here we use network analyses to identify the presence of co-flowering modules across seasons and quantify the phylogenetic diversity within each module. To do this, we tracked flowering phenology for plant communities at the Rocky Mountain Biological Laboratory (RMBL) in the central Colorado Rocky Mountains. The flowering species in this region are some of the best studied globally for flowering phenology (Dunne et al. 2003; Inouye 2008; Forrest et al. 2010; Aldridge et al. 2011; CaraDonna et al. 2014; Powers et al. 2022; Rivest et al. 2023) and plant-pollinator interactions (Bosch and Waser 1999; Forrest and Thomson 2010; Brosi and Briggs 2013; Waser and Price 2016; Gallagher and Campbell 2017; Ogilvie et al. 2017; Bain et al. 2022; Faust and Iler 2022; Schiffer et al. 2023). Different combinations of co-flowering species within these communities have been observed to either facilitate or interfere with pollination of one another (Forrest et al. 2010; Bain et al. 2022; Faust and Iler 2022; Arrowsmith et al. 2023; Schiffer et al. 2023), though the phylogenetic context of these interactions has not been explored.

Two previous studies have explored aspects of phylogenetic relationships in RMBL plant communities, to our knowledge. CaraDonna and Inouye (2015) tested for phylogenetic signal in multiple phenological traits of 60 species at RMBL, asking whether phenological traits tended to be more similar for more closely related species. They found significant phylogenetic signal in peak flowering and first flowering dates, but not in end date, flowering duration, or the sensitivities of any phenological trait to environmental variation (CaraDonna and Inouye 2015). Thus, the traits that will shape flowering overlap between these species include a mix of traits that are likely to follow patterns of phylogenetic relationships and those that will not. In a study of community phylogenetic diversity across elevations in the RMBL area, Bryant and colleagues (2008) found that plant communities tended to be increasingly overdispersed (more diverse than expected at random) as elevation increased up to observations at 3800 m above sea level, with a peak in absolute phylogenetic diversity circa 3000 m elevation, indicating that elevational gradients are likely important for shaping plant community diversity in this system.

Our study of the phenology of phylogenetic diversity in co-flowering communities includes wildflower communities in three meadows that range from a montane site at 2815 m above sea level to a subalpine site just below treeline at 3380 m. We quantify the distribution of flowering abundance of species in these communities and we use a network analysis to test whether flowering phenology is clustered into groups (modules) of co-flowering species across the season. We then quantify phylogenetic diversity across the season, and test whether it differs across modules of species flowering together. We test the general prediction that phylogenetic diversity will be higher than expected at these local scales and within co-flowering modules (overdispersed) but relatively lower (more underdispersed) early and late in the season when environmental conditions are most marginal. Finally, we ask whether overdispersion increases with elevation in co-flowering communities, as has been previously observed for co-occurring plants along the same gradient (Bryant et al. 2008).

## Materials & Methods

### Study Sites

We collected data at three sites in Washington Gulch near the Rocky Mountain Biological Laboratory (RMBL, Gothic, Colorado, USA) from June to August in 2021 and 2022. RMBL is in the East River valley of the West Elk mountains, approximately 10 kilometers from Crested Butte, Colorado. The relatively short growing season lasts 3-5 months, and snowmelt date is variable, but has been arriving earlier on average with climate change (Aldridge et al. 2011; CaraDonna et al. 2014). Study sites were located at 2815 m (38°53’50“N, 106°58’43”W), 3165 m (38°57’38“N, 107°01’53”W), and 3380 m (38°58’10“N, 107°01’53”W) in elevation.

These sites are part of a larger, long-term study of plant functional traits over an elevational gradient at RMBL established in 2003 (Bryant et al. 2008; Sloat et al. 2015). All sites are approximately 50 m^2^ and have similar slopes and aspects (Sloat et al. 2015). Using quadrats, we defined five 1.2 m x 1.2 m plots spaced 5 m apart from each other along a subtransect running downslope within each site. We did not control for density or other factors when placing the plots so we could capture an unbiased sample of the sites. We then marked the plots with flags at the corners and strings around the edges. We removed these markings at the end of each summer, so exact plot locations shifted slightly each year. The lowest and middle sites are in the montane vegetation zone, while the highest is subalpine (Ackerfield 2015). Across our sites, both soil pH and average July soil temperature decrease with elevation (Bryant et al. 2008). We identified forbs in the plots using the *Flora of Colorado* (Ackerfield 2015) and reference specimens at the Rocky Mountain Biological Laboratory Herbarium (herbarium code RMBL).

### Field Data Collection

Within the plots, we recorded all newly open flowers or inflorescences once per week from early June to mid-August for ten weeks of observation over each of the two summers. We counted flowers for all forbs in all plots, which resulted in counts for 56 species (see also Results). We counted flowers as ‘flowering units’, which we defined as either a single flower (most species), a capitula (Asteraceae), or an umbel (Apiaceae). We considered flowers open once the petals were separated and able to be counted. Each week, we tagged all new flowering units with a unique color of yarn and recorded the number of flowering units per species. We did not have predefined sample sizes and included all flowering units, which resulted in 3,660 flowering units in 2021, 1,973 in 2022, and a total of 5,633 flowering units across all sites and both years for our sample size (see also Results; Fig. 1). We only counted flowering units once as they opened, so counts each week were not cumulative. Marked flowering units were very rarely still open by the time of data collection for the next week, so recounting of previously-marked flowering units was not required to capture the flowering community each week (first author initials, pers. obs.). We never observed wilted or dead flowering units that we hadn’t previously tagged. All flowering units across all individuals of a species in all plots were combined to produce a single count of flowering units per species, per site, each week. Counts across weeks produced a distribution of flowering abundance across the 10 weeks of observation. This type of weekly abundance data has previously been shown to capture variation in co-flowering for co-flowering network analyses in similar-length growing seasons (Arceo-Gómez et al. 2018).

**Fig. 1:**
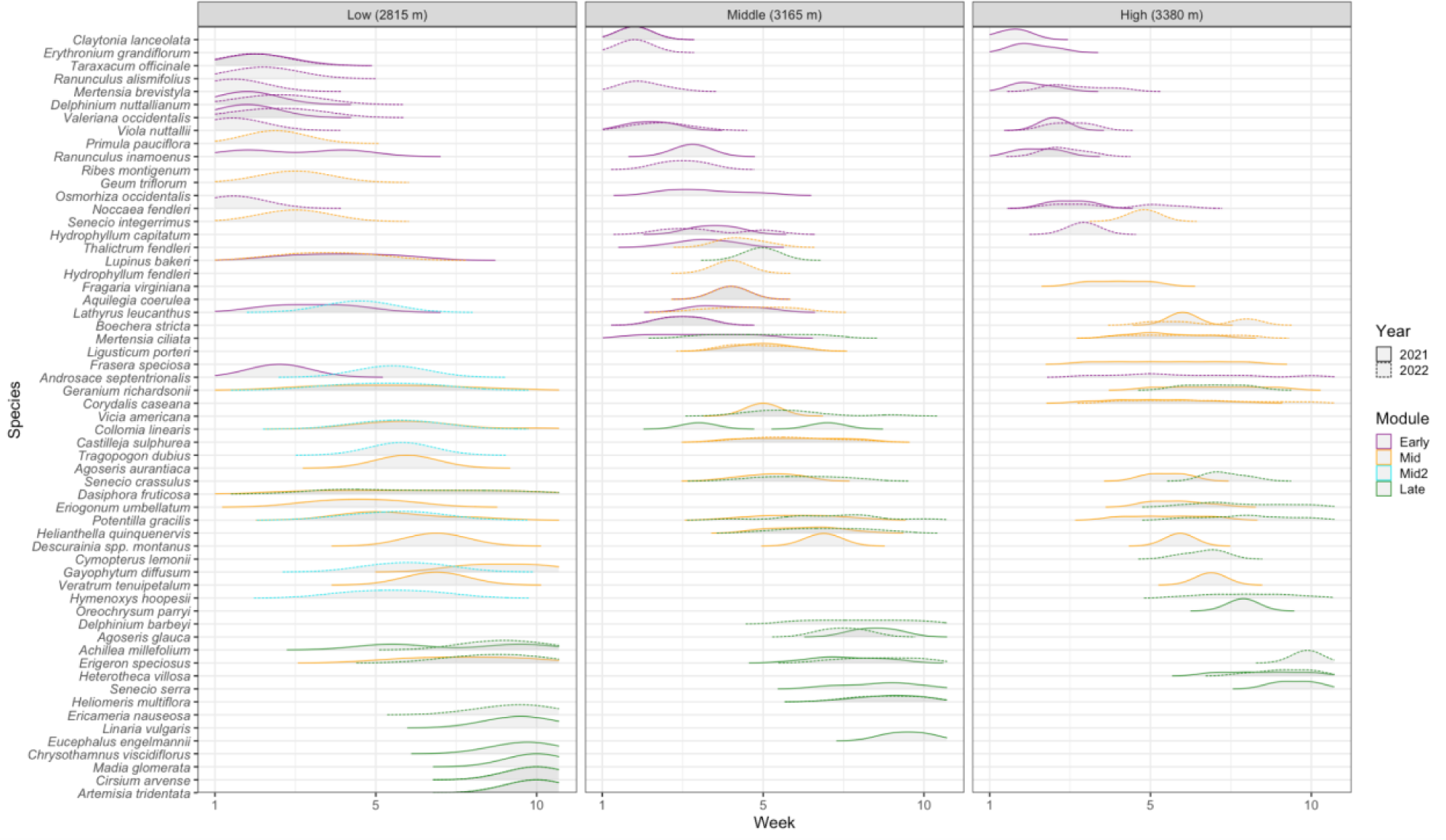
Density distributions for the number of flowering units (flowers or inflorescences) opened each week, for all species flowering at each site. Solid lines represent 2021 data and dashed lines represent 2022 data. Week 1 start dates were June 8, 2021 and June 6, 2022, and Week 10 start dates were August 9, 2021 and August 8, 2022. Lines are color coded by flowering module assigned by the network analysis (magenta for Early, orange for Mid, turquoise for a Mid2 module present only in the 2022 Low elevation site, and green for Late).

For species that fruited during the study period, we also quantified successful fruit and seed development at the end of each species’ flowering period. These data are not used for further analyses in this study, and data and associated Methods are available through the Environmental Data Initiative portal under package ID edi.1478.1.

### Identifying Modules in Co-flowering Networks

To test whether co-flowering communities were significantly modular, we used a network analysis with flowering species as the nodes and flowering overlap as the connections between them (edges). To quantify flowering overlap between each species pair in a given site and year, we calculated Schoener’s Index of niche overlap (Schoener 1970). We used the number of newly open flowers per week for each species to quantify its flowering phenology distribution and calculated Schoener’s Index between the distributions of a species pair as:

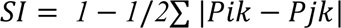

where *Pik* and *Pjk* are the proportion of species *i* and *j* flowering in week *k* (Forrest et al. 2010). Values range from 0 to 1, and fall at or near 0 when there is no flowering overlap and close to or at 1 for near or complete overlap in flowering distributions. We calculated Schoener’s Index in the R package spaa with the command niche.overlap() and method “schoener” (Zhang 2016).

We constructed co-flowering networks for each site for 2021 and 2022 (six total) with Schoener’s Index values as edge weights (Arceo-Gómez et al. 2018; Albor et al. 2020, 2022). An edge between two species will have more weight if they have a greater flowering overlap (i.e. a larger Schoener’s Index value). We constructed networks using the igraph R package (Csardi and Nepusz 2006). We calculated modularity using (Blondel et al. 2008)’s algorithm via the cluster.louvain() command in igraph (Table 1). Within the cluster.louvain() command, we specified weighted edges (according to their Schoener’s Index) and a resolution of 1.0. Resolution tells the algorithm whether to group the nodes into many, small modules or few, larger modules, and 1 is a standard value to not bias the algorithm towards module size (Newman and Girvan 2004; Csardi and Nepusz 2006).

**Table 1:**
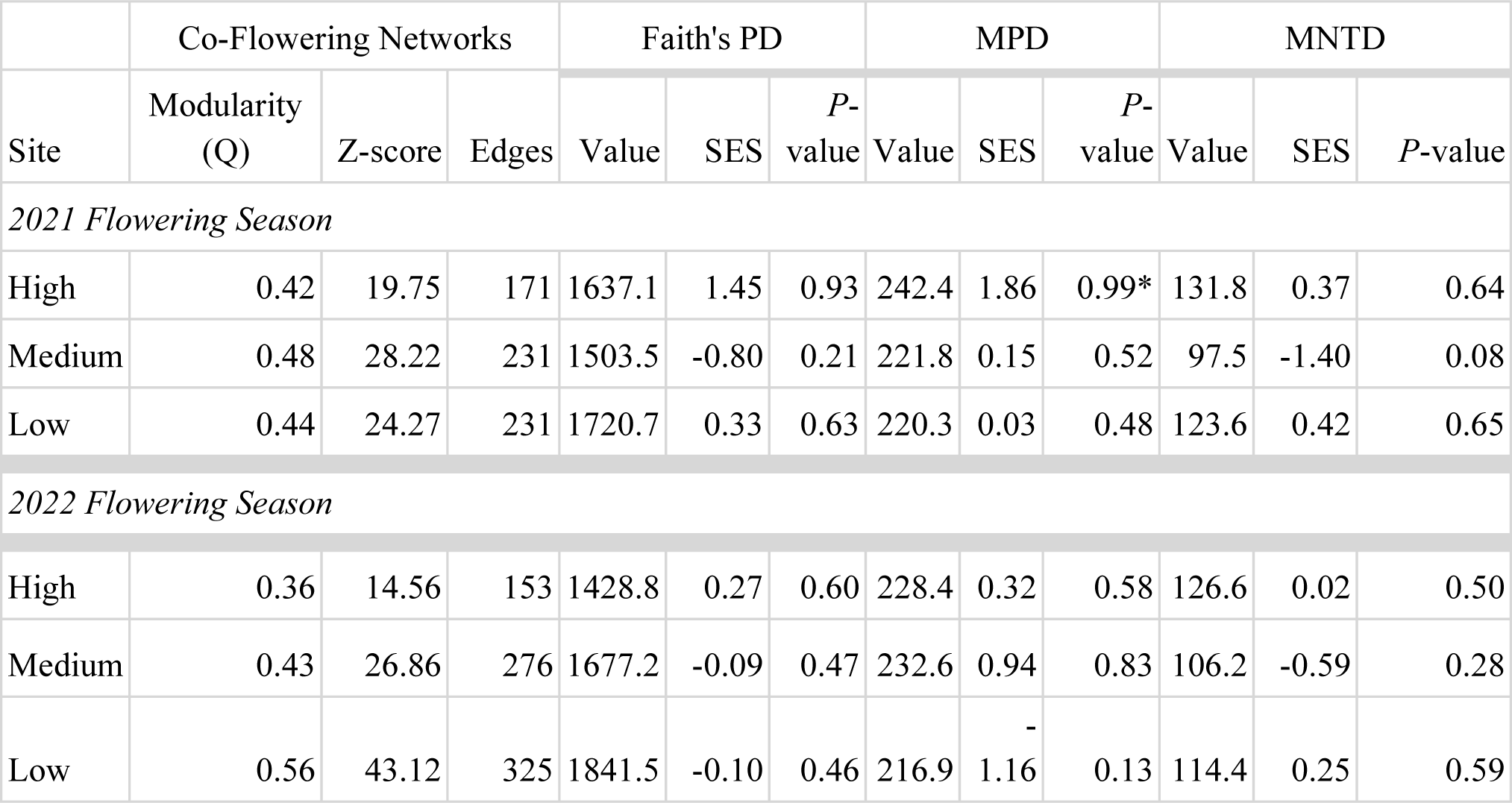
Metrics from the network and phylogenetic analyses for flowering communities at each site (High, Medium, Low) and year (2021, 2022). High site is at 3380 m, Medium is at 3165 m, and Low is at 2815 m. From the network analyses of co-flowering, shown are Modularity (Q scores), Z-score for modularity significance, and the number of Edges (species pairs) in each network. From analyses of phylogenetic diversity, shown are diversity metrics (Value), standard effect sizes (SES) for those values, and *P*-values for significance (for each phylogenetic diversity metric, including Faith’s Phylogenetic Diversity (PD), Mean Phylogenetic Diversity (MPD), and Mean Nearest Taxon Distance (MNTD). Significant P-values (those over above 0.975 or below 0.025) are indicated with an asterisk.

The modularity algorithm has two outputs: modularity values (Q), which range from −1 to 1, and which nodes (species) belong in which module. A modularity of Q = 1 represents a perfectly modular network in which modules have no connections (edges) outside their module. A Q = 0 indicates that the structure is no different than random, and Q = −1 results when there is no clustering in the network. To calculate the statistical significance of modularity (Q), we generated 1000 null networks for each of the six site and year combinations using the nullmodel() command and method “r2d” in the bipartite and vegan R packages (Dormann and Strauss 2014). We then calculated modularity (Q) of the null networks and a z-score for the observed modularity relative to the null distribution (Table 1). The z-score reflects how many standard deviations the actual network’s modularity is away from the mean modularity of all 1000 null networks, and z-scores above 2 are considered significant (Dormann and Strauss 2014).

We also used Gephi (Bastian et al. 2009) to calculate betweenness centrality, which is a network metric that estimates how many times a species connects otherwise unconnected species (ranging from zero to positive numbers of these connections), and can be used to identify keystone species with relatively high values that contribute uniquely to community overlap in flowering (Martín González et al. 2010; Arceo-Gómez et al. 2018). We constructed network visualizations in Gephi (Bastian et al. 2009).

### Phylogenetic Diversity of Co-Flowering Modules

To quantify and test predictions about phylogenetic diversity of co-flowering species across the season, we calculated phylogenetic diversity metrics for each site, and then for the communities of species flowering in each week and in each co-flowering module identified by the network analyses. We used the ‘ALLMB.tre’ angiosperm phylogeny from Smith and Brown (2018) to calculate all phylogenetic metrics. This phylogeny includes all genera in our dataset and 49 of 56 species observed in our plots. If a species was missing from the phylogeny but was the only member of its genus in the plots, we replaced it with a closely related congener in the Smith and Brown phylogeny (Supplementary material Table S1). Only the genus *Senecio* had multiple species in the plots missing from the phylogeny. We replaced *Senecio integerrimus* Nutt. with sister species *S. triangularis* Hook. (Pelser et al. 2007). We removed *Senecio crassulus* A. Gray from all phylogenetic analyses because it has no available genetic or phylogenetic information.

To quantify phylogenetic diversity, we calculated three metrics: Faith’s Phylogenetic Diversity (Faith 1992), Mean Phylogenetic Distance (MPD), and Mean Nearest Taxon Distance (MNTD) in the R package geiger (Pennell et al. 2014). Faith’s Phylogenetic Diversity is the total evolutionary distance (branch length) across a phylogenetic tree describing the relationships among the taxa in a dataset. This metric will increase if relationships are more distant across the tree (species are more evolutionarily divergent). Mean Phylogenetic Distance is the mean distance between pairs of species, and so it also increases as evolutionary relationships are more distant across the tree, but will be less sensitive to rare cases of very distantly-related species that would otherwise elevate Faith’s Phylogenetic Diversity. Mean Nearest Taxon Distance is the mean of the distance to the closest relative of each taxon, and so it reflects the typical distance to any species’ closest relative in the dataset and increases as closest relatives are more distant. Therefore, MNTD is more reflective of the tendency of species to have closer relatives than it is of the overall evolutionary diversity across the entire tree (community of species) that is captured by the former metrics (for further discussion, see review by (Tucker et al. 2017)).

To test whether a plot had higher phylogenetic diversity than expected (overdispersion) or less than expected (underdispersion), we calculated standard effect sizes (SES) for all three metrics in the picante R package (Kembel et al. 2010) using the ses.metric() function. Positive SES values indicate that the community in question is more phylogenetically diverse than a random draw of the same number of species from the regional species pool (overdispersion), and a *P* value above 0.975 indicates statistical significance of the SES. Negative SES values indicate that the species group in question is less phylogenetically diverse than a random draw of the same number of species from the regional species pool (underdispersion), and a *P* value below 0.025 indicates statistical significance of the SES. *P* values are significant if the SES absolute value is greater than 1.96. We used each metric’s “sample.pool” null model to draw a random sample of the same number of species as the focal community from the pool of all species observed across all RMBL sites with 5000 iterations.

We analyzed all data in R version 4.2.3 (R Core Team 2023).

## Results

### Flowering Observations

We recorded data for 56 species and 5,633 flowering units across all three sites and years (Fig. 1). We counted 3,660 flowering units in 2021 and 1,973 in 2022. Across both years, the lowest site had 33 species and 1,994 flowering units, the middle site had 29 species and 2,108 flowering units, and the highest site had 26 species and 1,531 flowering units. Five species occurred at all three sites: *Potentilla gracilis* Douglas ex Hook., *Viola nuttallii* Pursh, *Ranunculus inamoenus* Greene, *Mertensia brevistyla* S. Watson, and *Erigeron speciosus* DC (Fig. 1).

Our dataset of pairwise observations of flowering overlap within sites and years yielded a total of 1,262 pairwise observations, with site-year combinations ranging from a low of 136 pairs at the High site in 2022 to a high of 300 pairs at the Low site in 2022 (Supplemental File Schoeners_all.csv). Over half (57.2%) of the 1,262 species pair observations across sites and years had low or no overlap in their flowering phenology (Schoener’s Index at or near 0; Figs. S1 & S2). Over three quarters (76.4%) of these species pairs fell below a Schoener’s Index of 0.25 (lower quarter of the index, which ranges from 0 to 1). A small subset of species pairs had high flowering overlap: 58 of the 1,262 pairs (<5%) had Schoener’s Index values in the upper quartile of the index (at or above 0.75).

### Co-Flowering Network Modularity

We constructed networks for flowering communities at each site in each year. Network size ranged from 19 to 26 nodes (species) per network and from 153 to 325 edges (Schoener’s Index of overlapping flowering between species) each (Table 1; Fig. 2). All sites in both years were significantly modular (Q > 0.35 for all; Z-scores > 2 for all; Table 1), indicating that certain groups of species overlapped in their flowering more than with the other species outside of their module (Fig. 2). The High site had the lowest modularity (2021: Q= 0.42; 2022: Q=0.36) and the Low site had higher modularity (2021: Q=0.44; 2022: Q=0.56) in both years (Table 1), indicating tighter clustering of the modules (weaker overlap with other modules) in the Low site. All three sites had three modules each in 2021 (Fig. 2). In 2022, the High and Middle sites again had three modules, while the Low site had four (Fig. 2). The duration of flowering of species within each module varied, but ranged from 2-4 weeks, depending on the site, year, and time of the summer (Fig. 1).

**Fig. 2:**
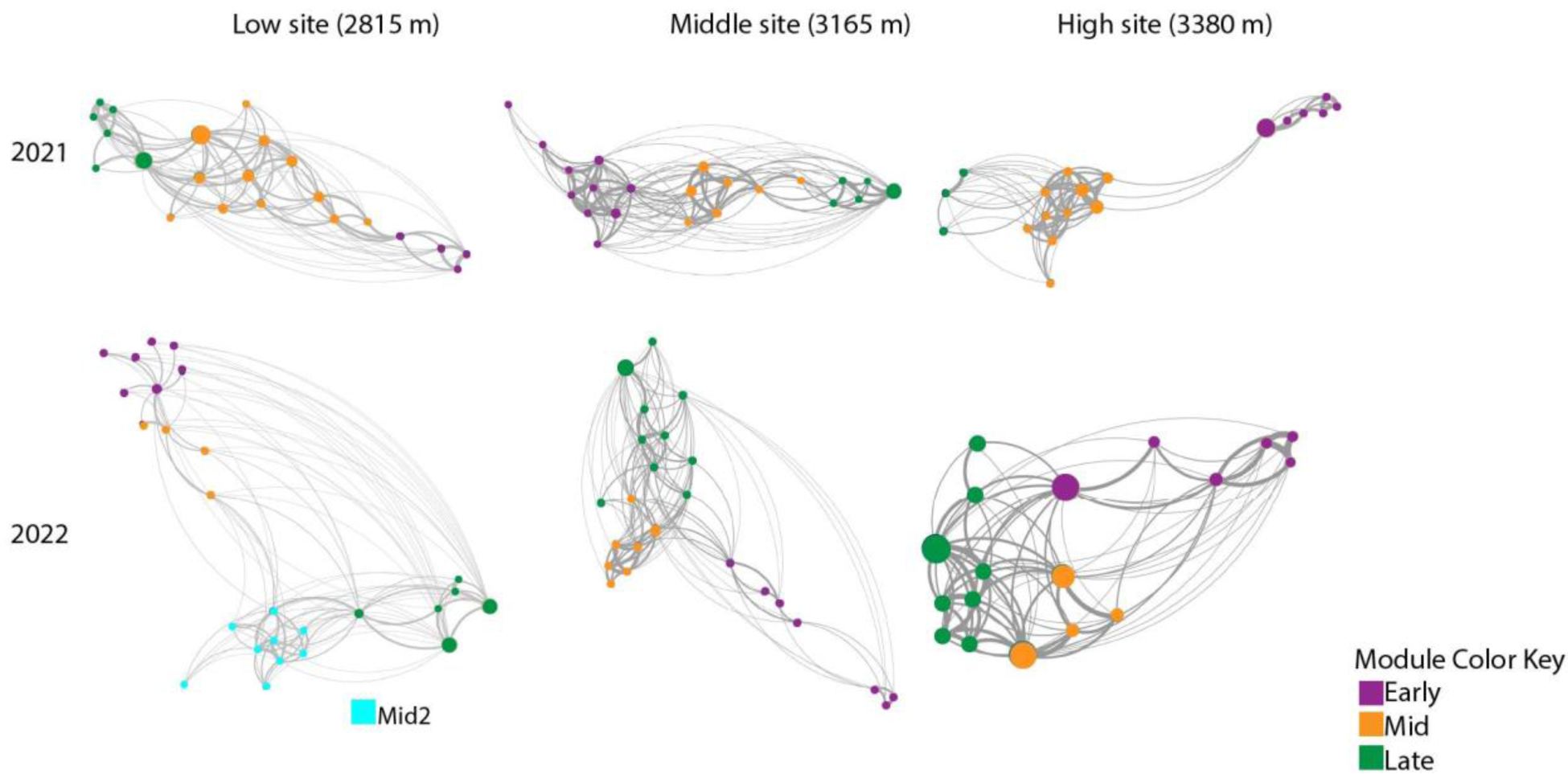
Network analyses of flowering time for species at each site and year, with nodes (species) color-coded according to the co-flowering module assigned by the analysis. Modules shown in purple flower early in the summer, followed by orange modules in the middle of the summer and green modules late in the summer. Node size represents betweenness centrality from the network analysis, with larger nodes showing species that co-flower with more species. Lines connecting species show edges weighted according to their Schoener’s Index of flowering overlap, where thicker lines have higher values (more flowering overlap) and thinner lines have lower values (less flowering overlap).

For all sites and years, modules corresponded with flowering at the early, mid, and late growing season, and approximate dates of the modules were relatively consistent for sites across both years (Fig. 1). A species’ module depended on when the largest proportion of its flowers were open. For example, if 90% of a species’ flowers were open in weeks 1 and 2, but 10% of its flowers opened in weeks 4 and 5, the modularity algorithm would place it in the early season module. Thus, it is possible for species flowering in the same week to be in different modules, depending on the other weeks when they had open flowers (Fig. 1). For the low site with an additional module in 2022, there were two middle-season modules spanning the middle of the growing season in July, but one had more overlapping flowering with the early season module and the other was less connected to modules before or after it (Figs. 1 & 2).

Betweenness centrality values indicated that a small set of species had large numbers of connections to otherwise unconnected species. The highest value of this metric was 197.03 for *Viola nuttallii* at the low elevation site in 2022 (Table S2). The average betweenness centrality value across our dataset was 8.6. In 2021, the average betweenness centrality was 5.6, and highest values were >30, seen for *Achillea millefolium, Dasiphora fruticosa, Heliomeris multiflora* and *Noccaea fendleri*. In 2022, the average betweenness centrality was 11.44, and highest values were >50, seen for *Tragopogon dubius, Artemisia tridentata, Lupinus bakeri, Erigeron speciosus, Ericameria nauseosa* and *Viola nuttallii*.

### Phylogenetic Diversity of Co-Flowering Modules

The 56 species in our plots spanned 24 families: 12 at the lowest site, 14 at the middle, and 16 at the highest. Five families (Asteraceae, Ranunculaceae, Rosaceae, Violaceae, Boraginaceae) occurred at all three sites (Fig. 3). At the site level, across the entire season, the Low site had the highest PD and lowest MPD in both years (Table 1). This indicates that the Low site tended to have lower diversity (per MPD) but had some divergent taxa elevating PD (Table 1). The only significant deviation from expected occurred for the High site in 2021, when it had significantly higher diversity than expected (overdispersion; Table 1). MNTD values were also the highest at the High site in both years, consistent with higher divergence between each flowering species and its closest relatives at this site (Table 1).

**Fig. 3:**
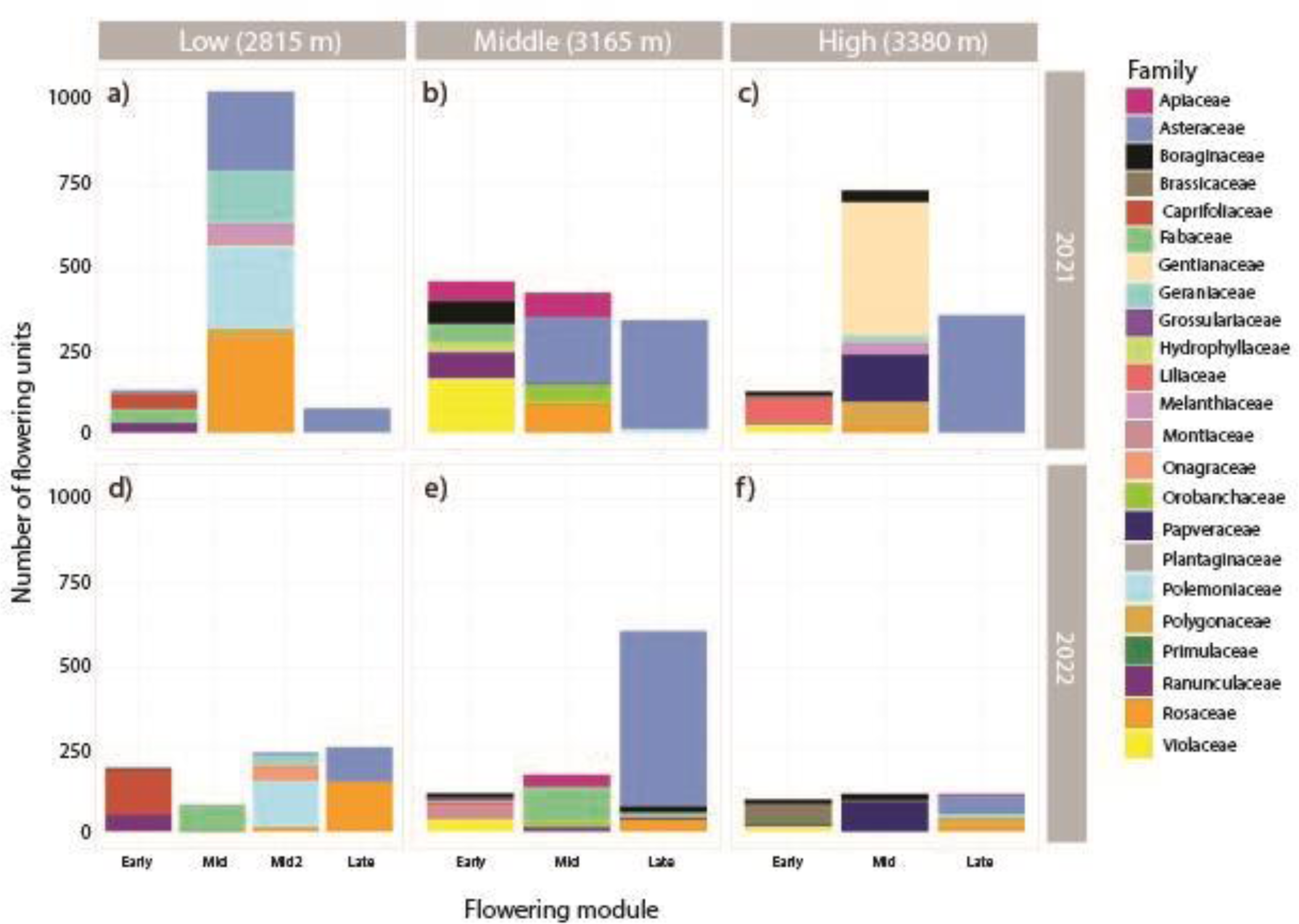
Histograms of the number of flowering units (flowers or inflorescences) color coded by family. Shown are panels for the Low elevation site (a, d), the Middle elevation site (b, e) and the High elevation site (c, f) for 2021 (a, b, c) and 2022 (d, e, f) flowering seasons. Each season is partitioned into co-flowering modules from the network analysis (Early, Mid, and Late, with an additional mid-season module in the 2022 Low elevation site).

Across weeks, phylogenetic diversity metrics generally indicated weak overdispersion (higher diversity than expected, but not significantly) before declining to underdispersion (lower diversity than expected) by the end of the season (Fig. 4). In 2021, underdispersion was significant for all metrics by Week 10 for both Middle and High sites, and for MPD at the Low site (Fig. 4). In 2022, underdispersion was weaker for the Middle and High sites in the later weeks, but highly significant for the Low site for all three metrics for the last three weeks of the season (Fig. 4).

**Fig. 4:**
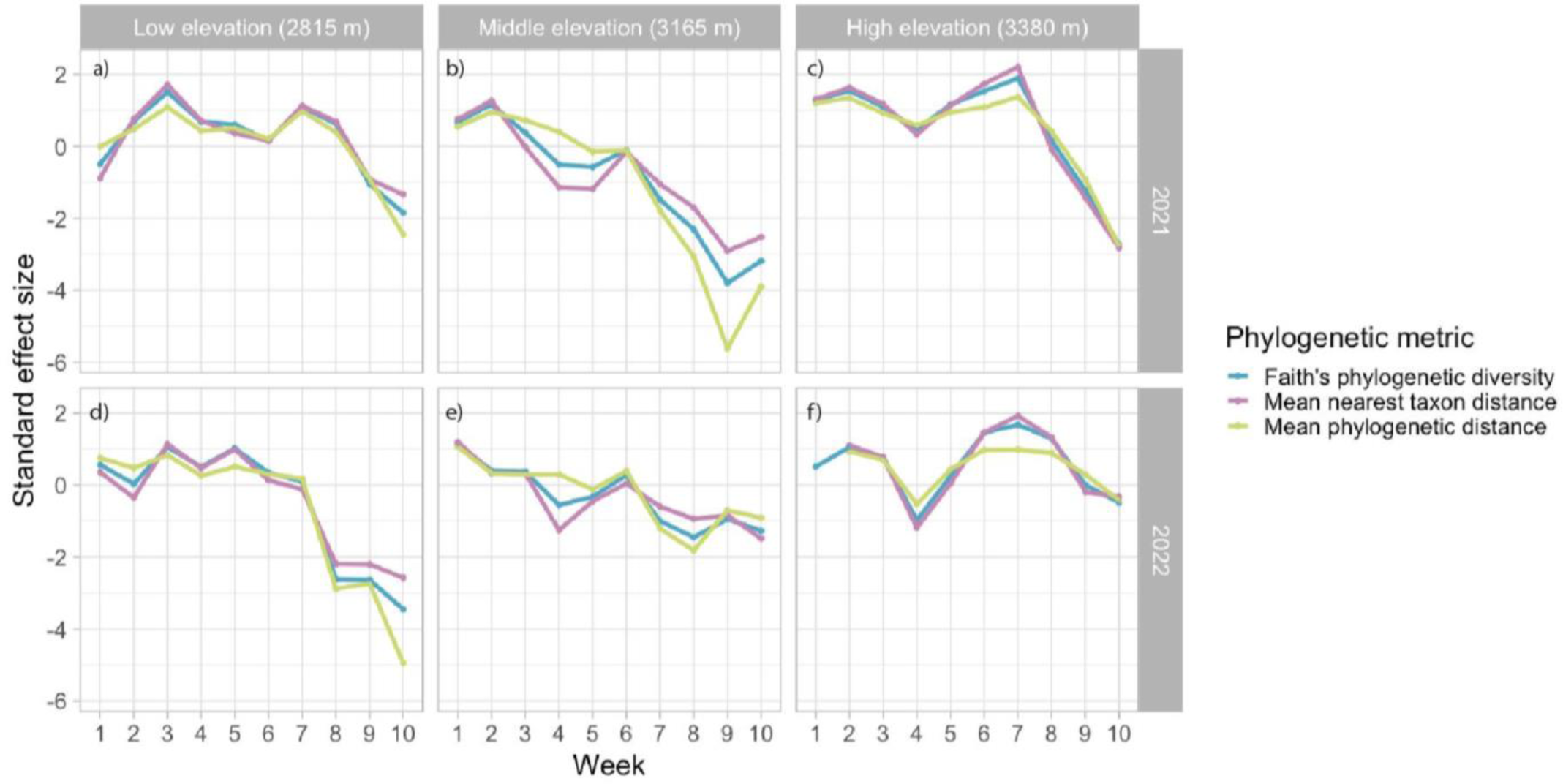
Standard effect sizes for metrics of phylogenetic diversity of the flowering community by week, for all sites and years. Metrics include mean nearest taxon distance (pink), mean phylogenetic distance (green), and Faith’s phylogenetic diversity (blue). Shown are panels for the Low elevation site (a, d), the Middle elevation site (b, e), and the High elevation site (c, f) for 2021 (a-c) and 2022 (d-f) flowering seasons. SES values of 1.96 or greater indicate significant overdispersion (greater diversity than expected), while values of −1.96 and below indicate significant underdispersion (lower diversity than expected).

For co-flowering modules of species flowering together, early-season modules were more overdispersed than late-season modules for all sites, years, and phylogenetic metrics (Fig. 5). Notably, all three metrics indicated significant underdispersion (SES < −1.96) for the late season module in 2021, and MPD was consistently lowest and strongly significant with an SES less than −4. This indicates that species co-flowering in the late season module are consistently more closely related to one another. In 2022, as for the weekly data, the tendency for overdispersion in the late season was present but weaker for the Middle and High sites, but continued to be highly significant for the Low site.

**Fig. 5:**
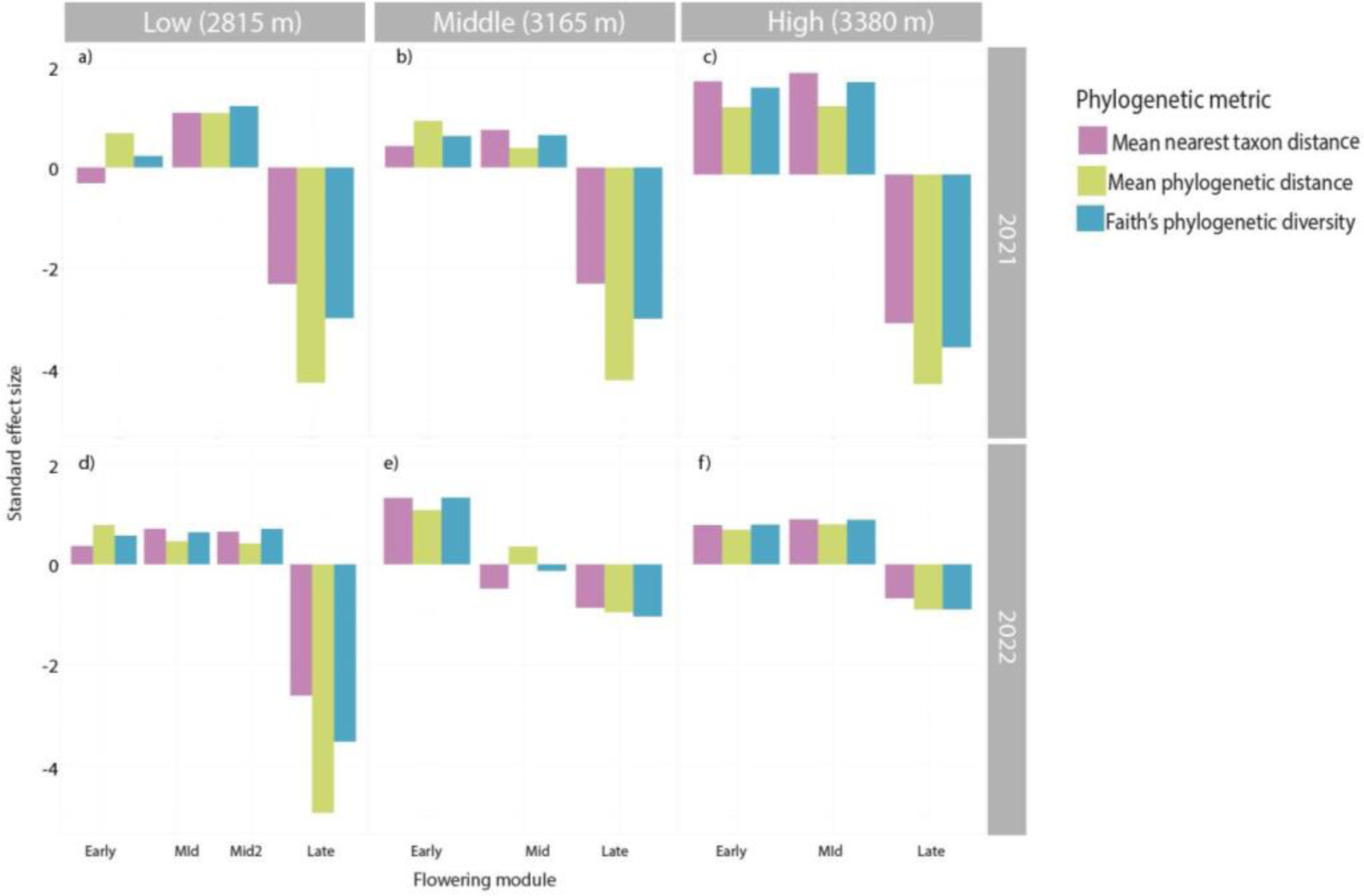
Standard effect sizes for metrics of phylogenetic diversity of species in co-flowering modules identified in the network analyses, for all sites and years. Metrics include mean nearest taxon distance (pink), mean phylogenetic distance (green), and phylogenetic diversity (blue). Shown are panels for the Low elevation site (a, d), the Middle elevation site (b, e), and the High elevation site (c, f) for 2021 (a, b, c) and 2022 (d, e, f) flowering seasons. Within sites and years, panels show phylogenetic diversity for each co-flowering module, including those flowering in early, mid, and late season. SES values of 1.96 or greater indicate significant overdispersion (greater diversity than expected), while values of −1.96 and below indicate significant underdispersion (lower diversity than expected).

## Discussion

Despite the importance of flowering phenology for plants’ reproductive success (Elzinga et al. 2007; Forrest and Miller-Rushing 2010) and persistence during environmental change (Cleland et al. 2007; Anderson 2016), the distribution of phylogenetic diversity across flowering seasons has been largely undescribed. Here, we find that Rocky Mountain wildflower communities are structured into modules of species that flower together, and that phylogenetic diversity varies among modules. Co-flowering modules tend towards phylogenetically overdispersion early in the season and at higher elevation, but become strongly underdispersed and depauperate in diversity at the end of the flowering season at all elevations.

Co-flowering was significantly modular at all three sites and in both years. The total richness of flowering taxa was approximately 30 species at each site (ranging from 33 at low elevation to 26 at high elevation), but for nearly all site-year combinations this diversity was divided into three modules of co-flowering species across the season. At the low elevation site in 2022, co-flowering species were further divided into an additional mid-season module. These seasonal modules ranged from 15% to 55% of all species that were observed to flower across the season at a site. Co-flowering communities were consistent across years: only 15 species switched modules from 2021 to 2022 across all sites, despite 2021 having nearly twice the abundance of flowers, with earlier snowmelt and less rain than 2022 (Prather et al. 2023). Thus, communities of co-flowering species contained a fraction of the total flowering diversity at each site, and co-flowered as largely the same communities year to year.

We predicted that phylogenetic diversity would be higher than expected within co-flowering modules (overdispersed) but relatively lower (more underdispersed) early and late in the season when environmental conditions are most marginal. We did not find significant overdispersion within any module in either year, and instead we found that modules tended towards overdispersion only earlier in the season (though not significantly). Phylogenetic diversity did decline towards the end of the growing season by all phylogenetic metrics, resulting in significant underdispersion by the end of the season. This seasonal pattern was consistent whether species were grouped by flowering week or by module, with declines at the end of the season beginning at roughly the same time as the transition to the flowering of the late season module. This variation in phylogenetic composition across co-flowering modules was associated with clear differences in the representation of regional taxonomic diversity. In particular, species in the Asteraceae and, to a lesser extent, the Rosaceae dominated the modules at the end of the flowering season. The prevalence of these taxa at the end of the season may be familiar to field biologists (first author initials, pers. obs), but to our knowledge, lower phylogenetic diversity of later flowering modules has not been previously reported. One previous study at RMBL found phylogenetic signal (clustering) in first and peak flowering dates, but not in last flowering date or flowering duration of species (CaraDonna and Inouye 2015). Those previously reported patterns are consistent with our finding that the late flowering community is phylogenetically clustered into relatively fewer clades, and that it occurs as a distinct module of species that begin their flowering late in the season.

Our observation of lower diversity at the end of the season might reflect more marginal conditions that are suitable for only a subset of species. Soil organic carbon, soil respiration, and net ecosystem productivity all decline throughout the season in this region (Moyes and Bowling 2013; Sloat et al. 2015). Declines in each of these variables have been associated with decreasing plant diversity in other systems (Tilman et al. 1997; Johnson et al. 2008; Lehmann et al. 2020).

Air temperature is also known to increase from June to August at this site, and has been shown to impact flowering phenology, soil temperature, and pollinator behavior (Harte et al. 1995; Aldridge et al. 2011; Ogilvie et al. 2017; Cordes et al. 2020). In contrast, while low soil moisture has been observed to cause declines in species richness and phylogenetic diversity in other systems (Deng et al. 2016; Wagg et al. 2017; Serafini et al. 2019; Miao et al. 2022), it is unlikely to be a factor here. Soil moisture in our system is affected by snowmelt timing and precipitation from summer thunderstorms, and has significant interannual variability (Pederson et al. 2011; Waser and Price 2016; Campbell et al. 2019; Mooney et al. 2021), which would not explain our consistent results.

Elevational differences could also be informative regarding the relationship between abiotic factors and phylogenetic diversity. Our high-elevation site as a whole was more overdispersed across both years by two of three diversity metrics (MPD and PD), and consistently more overdispersed than our low-elevation site. The low elevation site deviated little from null expectations and varied from weakly over-to underdispersed across metrics and years. A previous study by Bryant et al. (2008) across this same elevational gradient also found that phylogenetic overdispersion of plants increased with elevation, and that this relationship was predicted by lower soil temperature, lower pH, and higher total nitrogen as elevation increased. Similarly, Wolowski and colleagues found lower diversity and potential evidence of environmental filtering in lowland relative to montane sites for hummingbird-pollinated plants in the Brazilian Atlantic rainforest (Wolowski et al. 2017). These findings and ours suggest a pattern of lower phylogenetic diversity (underdispersion) of the flowering communities that occur in the warmer conditions at lower elevations and at the end of the season, and direct tests of this relationship would be valuable.

Studies in other systems have supported the opposite of our finding: underdispersion at high elevations, associated with abiotic filtering, wherein only a subset of evolutionary lineages share adaptations to the high elevation environment (Webb et al. 2002; Graham and Fine 2008; Dehling et al. 2014). These contrasting results highlight the uncertainty around which abiotic conditions are likely to impose phylogenetic filtering in different systems. We find evidence of filtering of flowering species only at the end of the flowering season, and thus the cooler/wetter conditions that occur early in the season and at higher elevations might be more favorable (less of a filter) for the species flowering across our gradient. It is noteworthy that the overdispersion at our highest site was weaker in 2022, when this region experienced a later snowmelt and wetter and cooler conditions (Prather et al. 2023), which may have imposed a stronger phylogenetic filter on our highest elevation site.

In addition to the effects of abiotic conditions, biotic interactions among co-flowering plants and their pollinators are likely to be shaping the diversity of flowering species across the season. Co-flowering has been studied as a mechanism of both and competition among plants, often specifically in the context of pollination (Sargent and Ackerly 2008; Carvalheiro et al. 2014; Briscoe Runquist et al. 2016). Pollinators can structure plant communities and co-flowering networks (Sargent and Ackerly 2008; Wei et al. 2021), and may impact co-flowering modules directly or indirectly through their interactions with abiotic conditions (Gallagher and Campbell 2017). Previous work at RMBL has found that pollen and pollinator limitation are often species-specific and/or context-dependent (Forrest and Thomson 2010; Gezon et al. 2016; Schiffer et al. 2023), which suggests that the mechanisms shaping co-flowering community assembly will depend on the particular species and context.

On the one hand, co-flowering is a potential mechanism of facilitation by which plants might overcome pollinator limitation and attract more pollinators to a multi-species patch (Ghazoul 2006). Indeed, some evidence suggests that co-flowering can increase pollinator visitation rates and the fitness of participating species by increasing the rewards for pollinators (Brown and Kodric-Brown 1979; Moeller 2004; Ghazoul 2006; Ogilvie and Thomson 2016; Bain et al. 2022). For example, in a recent study at RMBL, the experimental removal of the dominant plant species, *Helianthella quinquenervis,* immediately lowered pollinator visitation rates and network robustness for remaining species (Bain et al. 2022). In our network analysis, *H. quinquenervis* (Asteraceae) was observed as a middle or late season module species at the middle elevation site. It had betweenness centrality values that were lower than the average betweenness centrality of other species in both years, indicating low overlap with species outside of its co-flowering module and the potential for close interactions within its module, which would be dominated by related species in the Asteraceae in the late season. Similarly positive co-flowering interactions among related species have been seen in other systems. *Clarkia* congeners on the west coast of North America share pollinators and experience increased fitness and bee visitation through overlapping flowering (Moeller 2004; Moeller and Geber 2005*b*). Community members with similar floral shapes and sizes increased the seedset of *Delphinium barbeyi* at RMBL (Arrowsmith et al. 2023). Notably, facilitative interactions have been predicted and observed to be more common among distantly related species, which could result in overdispersion (Valiente-Banuet and Verdú 2007; Cavender-Bares et al. 2009), but the evidence above for pollinator facilitation among related species would predict that co-flowering modules would be more underdispersed.

On the other hand, sharing pollinators in time and space can lead to lower fitness through increased resource competition among plants (due to niche overlap) and/or reproductive interference (Levin and Anderson 1970; Johnson et al. 2022). These negative interactions could be stronger for closely related species if niches are phylogenetically conserved, though that need not be the case (Mayfield and Levine 2010). For example, work in the North American Great Lakes region showed that invasive *Lythrum salicaria* reduced bee visits to a native congener when they co-flowered (Brown et al. 2002). In a recent study at RMBL, Faust and Iler (2022) found that co-flowering decreased the seedset of *Linum lewisii*, but only when the plants were relatively less drought-stressed. Their results suggest that abiotic factors can mediate biotic co-flowering interactions, and the interactions may become more facilitative as environmental stresses increase (Faust and Iler 2022). That multiple interacting abiotic and biotic mechanisms might be shaping these co-flowering communities is suggested by the significant modularity in their networks. Arceo-Gomez et al. (2018) suggested that modularity in a network indicates multiple mechanisms affecting community assembly, because random or single mechanism-driven assembly would not result in clustering of groups of interacting species in time. Thus, future work should ideally combine tests of multiple abiotic and biotic interactions across phylogenetic divergence of community members to understand their assembly.

Finally, we note that flowering timing in these communities is already being affected by climate change. Globally, plants are flowering earlier with the advancing onset of spring (Menzel et al. 2006; Cleland et al. 2007; CaraDonna et al. 2014; Wolf et al. 2017), and this trend has been documented extensively at RMBL (Forrest et al. 2010; CaraDonna et al. 2014; Powers et al. 2022; Rivest et al. 2023). The patterns of co-flowering that we observe in this study are likely to be changing over time, and may not be optimized for current abiotic and biotic conditions. In particular, if climate change causes warm late season conditions to become more common earlier in the season (Inouye 2008; Pederson et al. 2011), then we might expect that the highest diversity early season modules are experiencing the most adverse effects. Studies of co-flowering network change over time would reveal whether co-flowering module communities shift their phenologies together, whether species diversity tends to be gained or lost in the process, and the potential roles of phylogenetic diversity in shaping response to environmental change.

In summary, we find that co-flowering communities are significantly modular, containing a subset of flowering species diversity at a site, and forming phenologically distinct communities that are largely consistent across years. We also find that late season modules have lower phylogenetic diversity according to all metrics (PD, MPD & MNTD) than early season modules, and that this decline in diversity occurred at all three elevations. These patterns are consistent with environmental filtering of phylogenetic diversity in the late season, but more work is needed to understand the abiotic and biotic conditions that might impose filtering on flowering of species at these sites. Across sites, our high elevation site contained more phylogenetic diversity than the low elevation site in both years, consistent with previous observations of overdispersion of the entire plant community at higher elevations, but contrasting with the prediction that high elevations should impose intense filtering and contain lower phylogenetic diversity. Our results clarify how phylogenetic diversity is distributed across sets of co-flowering species and across elevations in a subalpine plant community, but also highlight open questions about the mechanisms impacting the composition of co-flowering groups, and the evolutionary history of traits that allow for early or late season flowering.

