## Supplementary material for "Phylogenetic diversity of flowering plants declines across the growing season in Rocky Mountain wildflower communities"

Author names

**Supplementary Tables & Figures**

**Table S1:** Species in our research sites listed with replacement species used from Smith & Brown’s ALLMB.tre phylogeny (see also Methods). Road is the low elevation site at 2815 m, Pfeiler is the middle at 3165 m and PBM is the highest at 3380 m.

| RMBL species | Replacement from Smith & Brown 2018 | RMBL site(s) | Data year |
| --- | --- | --- | --- |
| *Senecio crassulus* | deleted | Pfeiler, PBM | 2022, 2021 |
| *Senecio integerrimus* | *Senecio triangularis* | Road, PBM | 2022 |
| *Cymopterus lemmonii* | *Cymopterus planosus* | PBM | 2022 |
| *Descurainia sp.* | *Descurainia incana* | PBM | 2021 |
| *Potentilla gracilis* | *Potentilla gracilis var. flabelliformis* | Pfeiler, PBM, Road | 2022, 2021 |
| *Veratrum tenuipetalum* | *Veratrum virginicum* | Road, PBM | 2021 |
| *Agoseris glauca* | *Agoseris glauca var. dasycephala* | Pfeiler | 2021 |
| *Eucephalus engelmannii* | *Eucephalus breweri* | Pfeiler | 2021 |
| *Helianthella quinquenervis* | *Helianthella uniflora* | Pfeiler | 2022, 2021 |
| *Hydrophyllum capitatum* | *Hydrophyllum capitatum var. capitatum* | PBM, Pfeiler | 2022, 2021 |
| *Gayophytum diffusum* | *Gayophytum diffusum subsp. diffusum* | Road | 2021 |
| *Madia glomerata* | *Madia sativa* | Road | 2021 |
| *Primula pauciflora* | *Primula pauciflora var. pauciflora* | Road | 2022 |

**Table S2**: Species in each network (site & year combination) listed with their betweenness centrality value. Average betweenness centrality is 8.6. Road is the low elevation site at 2815 m, Pfeiler is the middle at 3165 m and PBM is the highest at 3380 m.

| Species | Site | Year | Betweenness centrality |
| --- | --- | --- | --- |
| *Boechera stricta* | PBM | 2021 | 0 |
| *Claytonia lanceolata* | PBM | 2021 | 0 |
| *Oreochrysum parryi* | PBM | 2021 | 0 |
| *Pseudocymopterus montanus* | PBM | 2021 | 0 |
| *Veratrum tenuipetalum* | PBM | 2021 | 0 |
| *Viola nuttallii* | PBM | 2021 | 0 |
| *Senecio crassulus* | PBM | 2021 | 0.286 |
| *Senecio serra* | PBM | 2021 | 0.286 |
| *Heterotheca villosa* | PBM | 2021 | 0.943 |
| *Eriogonum umbellatum* | PBM | 2021 | 1.381 |
| *Mertensia ciliata* | PBM | 2021 | 1.381 |
| *Geranium richardsonii* | PBM | 2021 | 2.931 |
| *Potentilla gracilis* | PBM | 2021 | 2.931 |
| *Erythronium grandiflorum* | PBM | 2021 | 5 |
| *Mertensia brevistyla* | PBM | 2021 | 5 |
| *Ranunculus inamoenus* | PBM | 2021 | 5 |
| *Fragaria virginiana* | PBM | 2021 | 10 |
| *Corydalis caseana* | PBM | 2021 | 27.931 |
| *Frasera speciosa* | PBM | 2021 | 27.931 |
| *Noccaea fendleri* | PBM | 2021 | 65 |
| *Aquilegia coerulea* | Pfeiler | 2021 | 0 |
| *Claytonia lanceolata* | Pfeiler | 2021 | 0 |
| *Collomia linearis* | Pfeiler | 2021 | 0 |
| *Dasiphora fruticosa* | Pfeiler | 2021 | 0 |
| *Erythronium grandiflorum* | Pfeiler | 2021 | 0 |
| *Eucephalus engelmannii* | Pfeiler | 2021 | 0 |
| *Vicia americana* | Pfeiler | 2021 | 0 |
| *Boechera stricta* | Pfeiler | 2021 | 0.333 |
| *Hydrophyllum capitatum* | Pfeiler | 2021 | 0.7143 |
| *Ranunculus inamoenus* | Pfeiler | 2021 | 0.714 |
| *Agoseris glauca* | Pfeiler | 2021 | 0.75 |
| *Viola nuttallii* | Pfeiler | 2021 | 0.75 |
| *Senecio serra* | Pfeiler | 2021 | 1.917 |
| *Thalictrum fendleri* | Pfeiler | 2021 | 2.048 |
| *Ligusticum porteri* | Pfeiler | 2021 | 2.095 |
| *Senecio crassulus* | Pfeiler | 2021 | 2.095 |
| *Erigeron speciosus* | Pfeiler | 2021 | 4.05 |
| *Helianthella quinquenervis* | Pfeiler | 2021 | 4.162 |
| *Lathyrus leucanthus* | Pfeiler | 2021 | 7.798 |
| *Mertensia ciliata* | Pfeiler | 2021 | 7.798 |
| *Osmorhiza occidentalis* | Pfeiler | 2021 | 9.076 |
| *Castilleja sulphurea* | Pfeiler | 2021 | 9.495 |
| *Potentilla gracilis* | Pfeiler | 2021 | 9.495 |
| *Heliomeris multiflora* | Pfeiler | 2021 | 38.710 |
| *Agoseris aurantiaca* | Road | 2021 | 0 |
| *Androsace septentrionalis* | Road | 2021 | 0 |
| *Artemisia tridentata* | Road | 2021 | 0 |
| *Cirsium arvense* | Road | 2021 | 0 |
| *Delphinium nuttallianum* | Road | 2021 | 0 |
| *Gayophytum diffusum* | Road | 2021 | 0 |
| *Linaria vulgaris* | Road | 2021 | 0 |
| *Madia glomerata* | Road | 2021 | 0 |
| *Valeriana occidentalis* | Road | 2021 | 0 |
| *Veratrum tenuipetalum* | Road | 2021 | 0 |
| *Eriogonum umbellatum* | Road | 2021 | 0.571 |
| *Lathyrus leucanthus* | Road | 2021 | 0.761 |
| *Taraxacum officinale* | Road | 2021 | 1.209 |
| *Chrysothamnus viscidiflorus* | Road | 2021 | 1.55 |
| *Ranunculus inamoenus* | Road | 2021 | 1.752 |
| *Lupinus bakeri* | Road | 2021 | 5 |
| *Collomia linearis* | Road | 2021 | 9.321 |
| *Potentilla gracilis* | Road | 2021 | 9.733 |
| *Erigeron speciosus* | Road | 2021 | 10.549 |
| *Geranium richardsonii* | Road | 2021 | 10.983 |
| *Achillea millefolium* | Road | 2021 | 31.283 |
| *Dasiphora fruticosa* | Road | 2021 | 31.283 |
| *Erigeron speciosus* | PBM | 2022 | 0 |
| *Hydrophyllum capitatum* | PBM | 2022 | 0 |
| *Mertensia brevistyla* | PBM | 2022 | 0 |
| *Ranunculus inamoenus* | PBM | 2022 | 0 |
| *Senecio integerrimus* | PBM | 2022 | 0 |
| *Cymopterus lemmonii* | PBM | 2022 | 0.125 |
| *Eriogonum umbellatum* | PBM | 2022 | 0.125 |
| *Senecio crassulus* | PBM | 2022 | 0.286 |
| *Viola nuttallii* | PBM | 2022 | 0.333 |
| *Geranium richardsonii* | PBM | 2022 | 0.577 |
| *Hymenoxys hoopesii* | PBM | 2022 | 0.577 |
| *Heterotheca villosa* | PBM | 2022 | 1 |
| *Mertensia ciliata* | PBM | 2022 | 1.6 |
| *Noccaea fendleri* | PBM | 2022 | 1.933 |
| *Boechera stricta* | PBM | 2022 | 11.111 |
| *Androsace septentrionalis* | PBM | 2022 | 15.111 |
| *Corydalis caseana* | PBM | 2022 | 15.111 |
| *Potentilla gracilis* | PBM | 2022 | 15.111 |
| *Lupinus bakeri* | Pfeiler | 2022 | 0 |
| *Thalictrum fendleri* | Pfeiler | 2022 | 0 |
| *Ribes montigenum* | Pfeiler | 2022 | 0.2 |
| *Agoseris glauca* | Pfeiler | 2022 | 0.5 |
| *Heliomeris multiflora* | Pfeiler | 2022 | 0.5 |
| *Claytonia lanceolata* | Pfeiler | 2022 | 0.567 |
| *Erythronium grandiflorum* | Pfeiler | 2022 | 0.567 |
| *Mertensia brevistyla* | Pfeiler | 2022 | 1 |
| *Aquilegia coerulea* | Pfeiler | 2022 | 1.167 |
| *Hydrophyllum fendleri* | Pfeiler | 2022 | 1.1677 |
| *Delphinium barbeyi* | Pfeiler | 2022 | 1.397 |
| *Boechera stricta* | Pfeiler | 2022 | 1.4 |
| *Viola nuttallii* | Pfeiler | 2022 | 1.867 |
| *Castilleja sulphurea* | Pfeiler | 2022 | 2.167 |
| *Castilleja sulphurea* | Pfeiler | 2022 | 2.931 |
| *Mertensia ciliata* | Pfeiler | 2022 | 3.29 |
| *Senecio crassulus* | Pfeiler | 2022 | 3.29 |
| *Vicia americana* | Pfeiler | 2022 | 3.29 |
| *Helianthella quinquenervis* | Pfeiler | 2022 | 3.702 |
| *Ligusticum porteri* | Pfeiler | 2022 | 5.869 |
| *Hydrophyllum capitatum* | Pfeiler | 2022 | 14.433 |
| *Lathyrus leucanthus* | Pfeiler | 2022 | 16.202 |
| *Potentilla gracilis* | Pfeiler | 2022 | 18.156 |
| *Erigeron speciosus* | Pfeiler | 2022 | 66.34 |
| *Cirsium arvense* | Road | 2022 | 0 |
| *Collomia linearis* | Road | 2022 | 0 |
| *Delphinium nuttallianum* | Road | 2022 | 0 |
| *Erigeron speciosus* | Road | 2022 | 0 |
| *Gayophytum diffusum* | Road | 2022 | 0 |
| *Lathyrus leucanthus* | Road | 2022 | 0 |
| *Mertensia brevistyla* | Road | 2022 | 0 |
| *Noccaea fendleri* | Road | 2022 | 0 |
| *Potentilla gracilis* | Road | 2022 | 0 |
| *Primula pauciflora* | Road | 2022 | 0 |
| *Ranunculus alismifolius* | Road | 2022 | 0 |
| *Taraxacum officinale* | Road | 2022 | 1.2 |
| *Androsace septentrionalis* | Road | 2022 | 2.666 |
| *Dasiphora fruticosa* | Road | 2022 | 4.133 |
| *Senecio integerrimus* | Road | 2022 | 7.633 |
| *Valeriana occidentalis* | Road | 2022 | 7.633 |
| *Geranium richardsonii* | Road | 2022 | 10.5 |
| *Hymenoxys hoopesii* | Road | 2022 | 10.5 |
| *Achillea millefolium* | Road | 2022 | 16 |
| *Geum triflorum* | Road | 2022 | 16.26 |
| *Tragopogon dubius* | Road | 2022 | 53.03 |
| *Artemisia tridentata* | Road | 2022 | 62.76 |
| *Lupinus bakeri* | Road | 2022 | 62.76 |
| *Ericameria nauseosa* | Road | 2022 | 101.86 |
| *Viola nuttallii* | Road | 2022 | 197.03 |

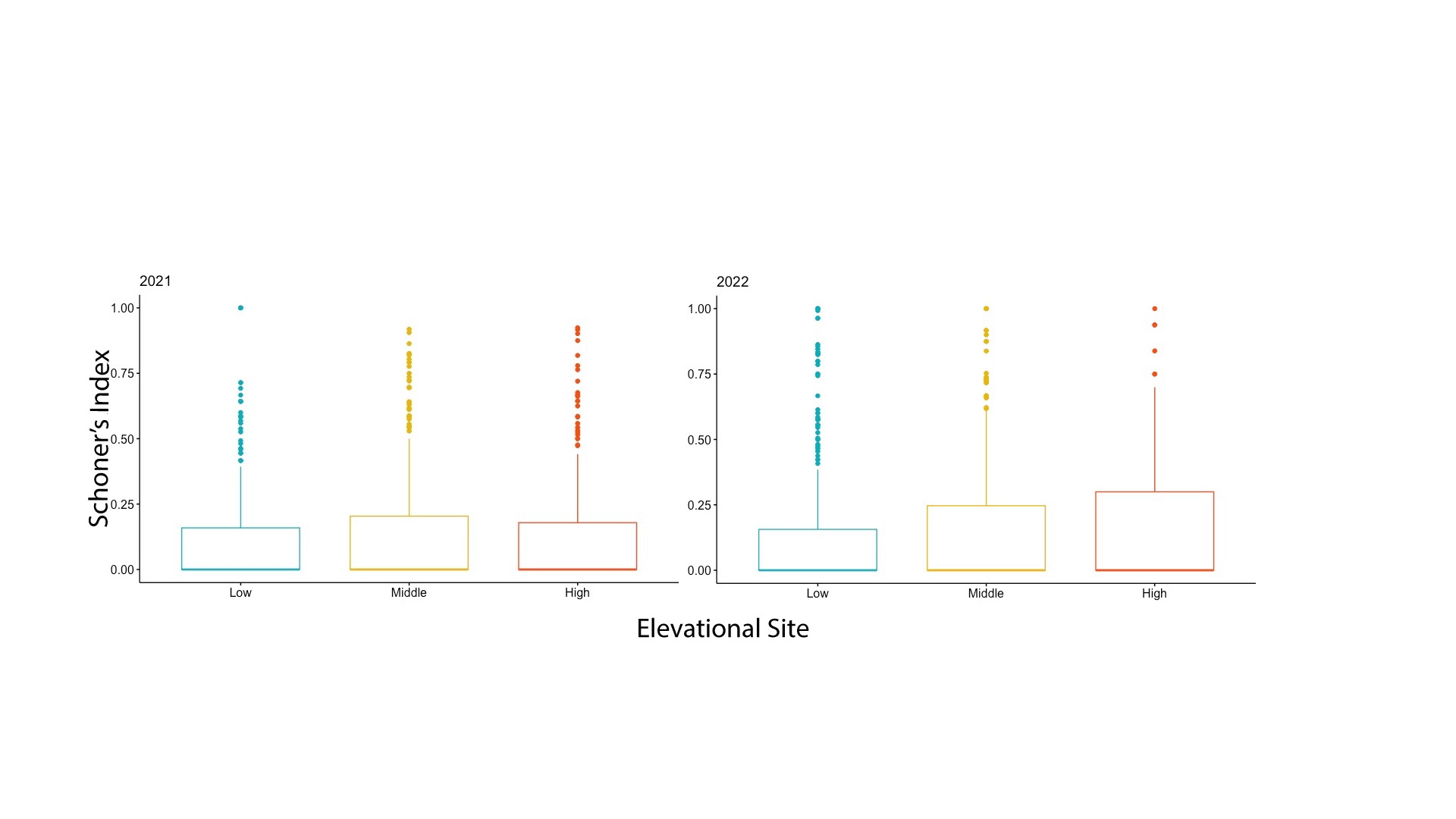

**Figure S1:** Box plots showing the distribution of Schoner’s Index values across sites for both years. Kruskal-Wallis tests for differences between the sites showed no significant difference in 2021 (χ2 = 6, df =2, p=0.06) and a significant difference in 2022 (χ2 = 13, df =2, p=0.002).

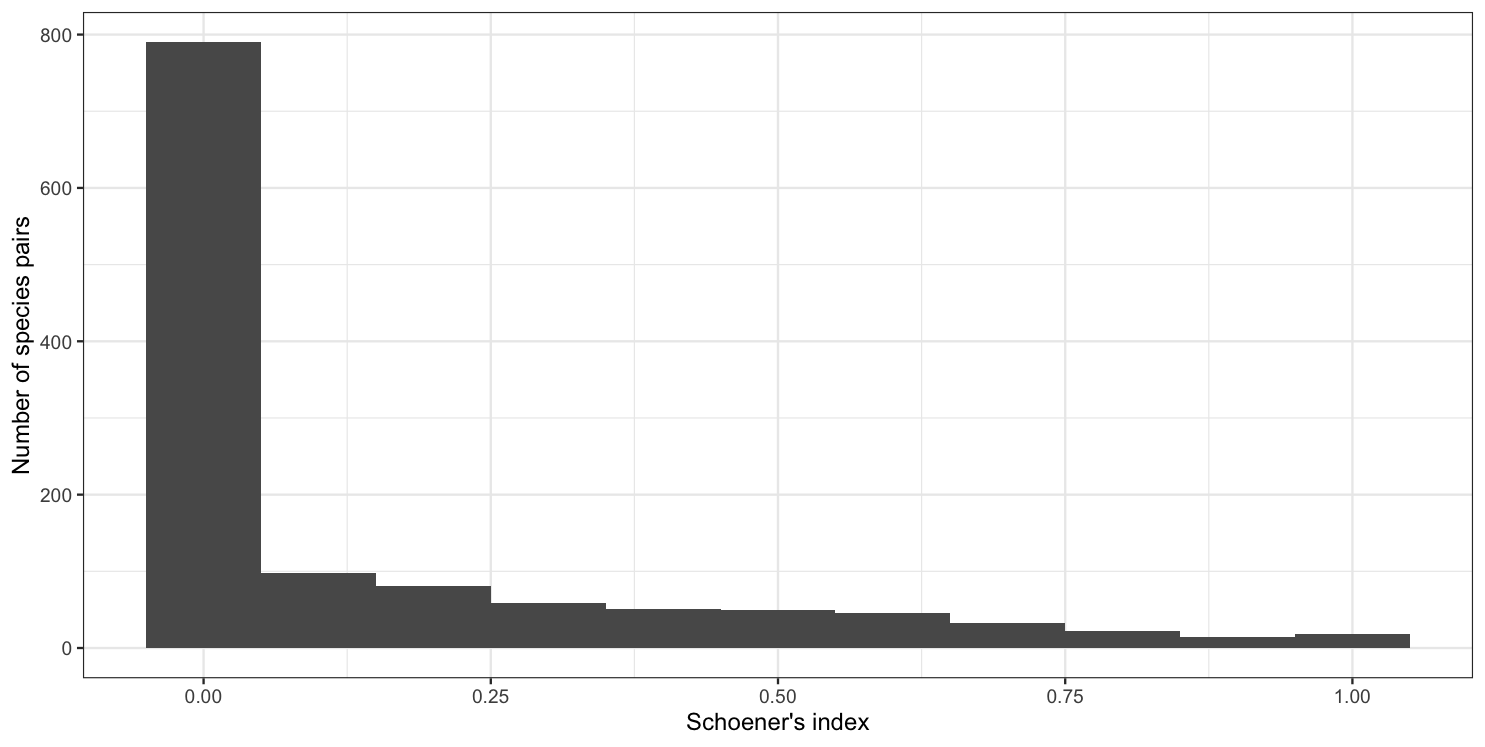

**Figure S2:** Histogram showing the distribution of Schoener’s Index values for all species pairs across all sites and years.
